## Supplementary Information for "Kinase KEY1 controls pyrenoid condensate size throughout the cell cycle by disrupting phase separation interactions"

### Supplemental information

**Supplementary Video 1. Pyrenoid condensates in wild type, *key1-1* mutant, and the rescued strain of *key1-1* (*key1-1;KEY1-SNAP*), related to Fig. 1.** Videos of confocal z-slices of overlay of RBCS1-Venus (green) and chlorophyll autofluorescence (magenta) through the entire cells shown in Fig. 1f-h. Left: wild type; middle: *key1-1* mutant; right: the rescued strain of *key1-1* (*key1-1;KEY1-SNAP*). Cells were unsynchronized and grown under constant light. Z-step = 300 nm.

**Supplementary Video 2. KEY1 is required for the dissolution of pyrenoid condensate (visualized by EPYC1-Venus) during cell division, related to Fig. 2.** Videos of the cell division shown in Fig. 2a,b,d,e. Top: overlay of the maximum intensity Z-projection EPYC1-Venus (green) and chlorophyll autofluorescence (magenta) for wild-type (left) and *key1-1* mutant (right) cells. The chlorophyll autofluorescence is displayed as the midplane Z-slice and maximum intensity Z-projection respectively. Bottom: heat map of the Venus channel alone for both cell types, with the scale identical to that in Fig. 2b,e. The first cell division occurs at 0 min for each cell.

**Supplementary Video 3. KEY1 is required for the dissolution of pyrenoid condensate (visualized by RBCS1-Venus) during cell division, related to Fig. 2.** Videos of maximum Z-projections of confocal timelapses of the cell divisions shown in Extended Data Fig. 2b,c,e,f. Top: overlay of the RBCS1-Venus (green) and chlorophyll autofluorescence (magenta) channels for wild-type (left) and *key1-1* mutant (right) cells. Bottom: heat map of the Venus channel alone, with the scale identical to that in Extended Data Fig. 2c,f. The first cell division occurs at 0 min for each cell.

**Supplementary Video 4. Pyrenoid condensate components' dissolution dynamics (visualized by RBCS1-mCherry and EPYC1-Venus) are highly correlated during cell division, related to Fig. 2.** Videos of maximum Z-projections of confocal timelapses of a representative wild-type cell division (left: chlorophyll autofluorescence, middle: RBCS1-mCherry, right: EPYC1-Venus). The first cell division occurs at 0 min.

**Supplementary Video 5. Ectopic pyrenoid condensate nucleated in the chloroplast in the *key1-1* mutant cell during cell growth at the beginning of the light cycle, related to Fig. 2.** Videos of confocal z-slices of an overlay of RBCS1-Venus (green) and chlorophyll autofluorescence (magenta) of *key1-1* mutant cell shown in Extended Data Fig. 3c,e for hourly time points. Hours indicate the time from the start of the light cycle. The white arrows indicate a *de novo* ectopic pyrenoid condensate. Z-step = 400 nm.

**Supplementary Video 6. Rotating view of the KEY1 3D structure predicted by AlphaFold 2, related to Fig. 5.** A video of a rotating ribbon diagram of AlphaFold 2-predicted structure of KEY1 shown in Fig. 5c. The Rubisco-binding motif is shown in black, the Uniprot-predicted disordered regions are shown in cyan, and the protein kinase domain is shown in orange.

**Supplementary Video 7. A simulation of KEY1-regulated pyrenoid condensate dynamics during cell division in a two-dimensional geometry mimicking the shape of the Chlamydomonas chloroplast, related to Fig. 6.** To recapitulate the hypothesized temporal changes in KEY1 activity during cell division, the EPYC1 phosphorylation rate was varied over time (left)**.** The resulting spatiotemporal phase transitions in a geometry that mimics the shape of the Chlamydomonas chloroplast are shown as the volume fraction of EPYC1 in green (right). The instantaneous phosphorylation rate is noted by a dot in the plot on the left.

**Supplementary Video 8. KEY1 activity drives size control of condensates in minimal model, related to Fig. 6.** The model was initialized with two EPYC1 condensates of different sizes at a constant intermediate phosphorylation rate; the rate is the same as in Fig. 6b,c iv. The resulting phase behaviors are shown by EPYC1 volume fraction in green. This video corresponds to Fig. 6d.

**Supplementary Video 9. KEY1 activity drives self-centering behavior of condensate in minimal model, related to Fig. 6.** The model was initialized with an off-center condensate with a constant low phosphorylation rate; the rate is lower than in Fig. 6d,f. The resulting phase behaviors are shown by EPYC1 volume fraction in green. This video corresponds to Fig. 6e.

**Supplementary Video 10. In the absence of KEY1 activity, condensates do not exhibit self-centering behavior in minimal model, related to Fig. 6.** The model was initialized with an off-center condensate and no kinase or phosphatase activity. The resulting phase behaviors are shown by EPYC1 volume fraction in green.

**Supplementary Video 11. KEY1 activity drives condensates repulsion at short distance, related to Fig. 6.** The model was initialized with two EPYC1 condensates that were close together. The resulting phase behaviors are shown by EPYC1 volume fraction in green. This video corresponds to Fig. 6f.

**Supplementary Video 12. Low KEY1 activity is sufficient to dissolve ectopic EPYC1 condensates, related to Fig. 6.** The model was initialized with a low phosphorylation rate and two EPYC1 condensates: one at the typical pyrenoid location and one elsewhere in the chloroplast geometry. The resulting phase behaviors are shown by EPYC1 volume fraction in green. This video corresponds to Fig. 6g (bottom).

**Supplementary Video 13. No KEY1 activity and increased self-attractive interactions leads to stable ectopic clusters.**
The model was initialized with no switching rates and two EPYC1 condensates: one at the typical pyrenoid location and one elsewhere in the chloroplast geometry. The resulting phase behaviors are shown by EPYC1 volume fraction in green. This video corresponds to Fig. 6g (top).

**Supplementary Video 14. Dynamic phase transitions in the minimal model are also seen in a square geometry with periodic boundary conditions, related to Fig. 6.**
The model was computed in the same manner as Supplementary Video 7 except in a square geometry with periodic boundary conditions. This video corresponds to Fig. 6b,c.
